# Cell-Type Switches Induced by Stochastic Histone Modification Inheritance

**DOI:** 10.1101/419481

**Authors:** Rongsheng Huang, Jinzhi Lei, Zhou Pei-Yuan

## Abstract

Cell plasticity is important for tissue developments during which somatic cells may switch between distinct states. Genetic networks to yield multistable states are usually required to yield multiple states, and either external stimuli or noise in gene expressions are trigger signals to induce cell-type switches between the states. In many biological systems, cells show highly plasticity and can switch between different state spontaneously, but maintaining the dynamic equilibrium of the cell population. Here, we considered a mechanism of spontaneous cell-type switches through the combination between gene regulation network and stochastic epigenetic state transitions. We presented a mathematical model that consists of a standard positive feedback loop with changes of histone modifications during with cell cycling. Based on the model, nucleosome state of an associated gene is a random process during cell cycling, and hence introduces an inherent noise to gene expression, which can automatically induce cell-type switches in cell cycling. Our model reveals a simple mechanism of spontaneous cell-type switches through a stochastic histone modification inheritance during cell cycle. This mechanism is inherent to the normal cell cycle process, and is independent to the external signals.

## 1. Introduction

Cell-type switches are important in mammalian development and the progress of of complex diseases, such as immune responses and drug resistance. At the molecular level, cell-type switch is often marked by variants of gene expression in marker genes, which show plasticity of cell types over time. The underling mechanisms to regulate the dynamical process of cell-type switch are important for our understanding of embryo developmental and disease progressing [62].

Cell type switches often associate with multiple expression modes of marker genes; switches between different cell types can be induced by either stochastic gene expression or external regulations in the sense of bifurcation [12, 15, 31, 33, 34,48,52,55,57,61]. Stochastic gene expression is a main source of phenotype switch in bacteria[7]; the stochasticity comes from both intrinsic noise of promotor activities and extrinsic noise from environmental fluctuations. In mammalian cells, changes in the activities of regulators in the gene regulation network is one of the driving force to induce cell type switch, *e.g.*, stem cell differentiation [10, 23, 24, 42]. During development, cells undergo a unidirectional course of differentiation, which can be viewed as a dynamical process on the Waddington epigenetic landscape with multistable states, the input signals can change the landscape to guide the transition between different states [15, 19, 72].

Chromatin regulators play crucial roles in establishing and maintaining gene expression states [4, 30]. Histone modifications and DNA methylations are important epigenetic states that can regulate the chromatin state of a DNA sequence, and modulate the gene expression dynamics. These epigenetic modifications are modulated by intercellular enzyme activities, and hence dynamically change over time and during cell cycle [50]. The stochastic epigenetic inheritance during cell cycle can result in spontaneous alteration of epigenetic state during development; epigenetic changes in regulatory loci often correlate with expression changes during stem cell differentiation [3, 8, 27, 58, 73].

Here, to investigate how stochastic epigenetic inheritance affect the dynamical process of cell fate decision, we studied a computational model that combines positive feedback regulation with histone modification of the gene promoter. Moreover, cell cycle was included in the model through the stochastic inheritance during cell cycle. We show that the dynamic behavior of histone modification in the cell division can induce spontaneous cell-type switches during normal development process, from which the Waddington landscape is constructed. Moreover, we show that the combination of feedback regulation with histone modification can produce intermediate cell states that are essential in EMT and cell plasticity.

## 2. Model

### 2.1. Model description

To investigate the role of histone modification in cell type switch, we considered a simple model that combine a self-activation positive feedback with stochastic histone modification in the promoter region (Fig. 1). We assumed that the cell type is represented by the expression of a marker gene; the encoded protein regulates (either directly or indirectly) its own promoter activity to form a positive feedback to maintain the bistable transcriptional landscape. This type of simple motif has been extensively studied in many systems with different states in response to various induction stimulus, variance in the induction stimulus are able to induce switches between the two stable states [1, 16, 22, 25, 41, 47, 64]. In this study, to study how nucleosome states affect the cell-type switch, we further assumed that the positive feedback is modulated by the histone modification of nucleosomes in the promoter region; the proteins active its own transcription by binding to the promoter region, and the binding affinity is dependent on the nucleosome state, which is stochastically change over cell cycle [50].

**Figure 1.**
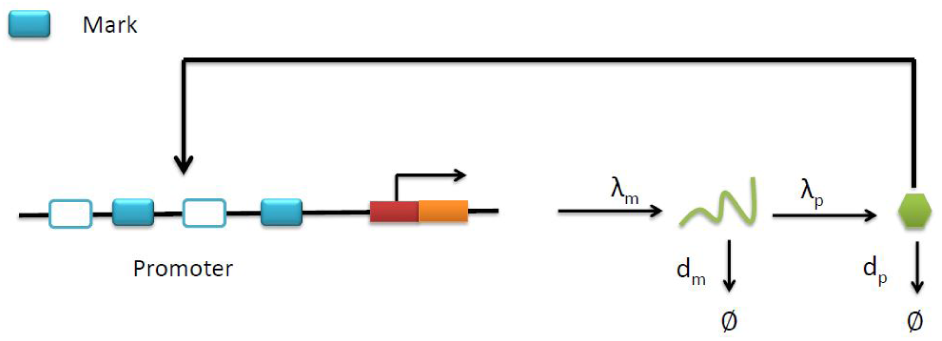
Illustration of gene regulation combines a positive feedback with stochastic histone modification at the enhancer region. Histone modifications alter the 50% effective concentration (EC50) of the toggle switch to regulate gene expression (see Formulations for details).

In eukaryotic cells, most DNA sequences are enclosed into basic organizational chromatin units–nucleosomes. A typical nucleosome has approximately 147 nucleotide base pairs that wrap around a histone octamer; the octamer is composed of one (H3 H4)_2_ tetramer capped by two H2A-H2B dimers[21]. The *N* -tail of the two copies of histone H3 can undergo various types of covalent modifications; the modification can either repress or active gene expression. For example, the trimethylation of H3 lysine 4 (H3K4me3) is often associated with active transcription, and the trimethylation of H3 lysine 27 (H3K27me3) is associated with transcription repression. During erythroblast differentiation, changes in the markers of H3K4me3 and/or H3k27me3 are seen in different genes [8, 58]. In our study, we focused at the gene expression activation modification H3K4me3, and assumed that the modification increases the promotion of the protein to its own transcription.

### 2.2. Formulations

Denote *m* and *p* respectively as the mRNA and protein concentrations of the marker gene. The proteins are translated from mRNA, and positive regulate the mRNA transcription. Hence, we have the following based differential equations:

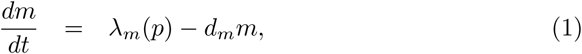

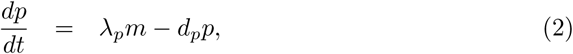

where *λ*_*m*_, *λ*_*p*_ are transcription and translation rates, and *d*_*m*_, *d*_*p*_ are degradation/dilution rates of mRNA and proteins. The transcription rate *λ*_*m*_ was assumed to be dependent on the protein concentration through a Hill type function[6, 17]:

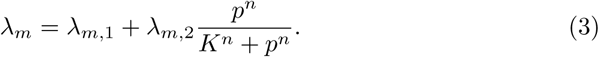

Here, *λ*_*m,*1_ is the basal transcription rate, and *λ*_*m,*2_ the maximum promotion in transcription due to the protein regulation, the coefficient *K* (50% effective concentration, EC50) measures the binding affinity of the protein to the enhancer, lower *K* measures higher affinity.

While we assumed that the nucleosome state affects the binding affinity through the modulation of chemical potential of the reactions, the EC50 *K* depends on the number of active markers *u* through an exponential function

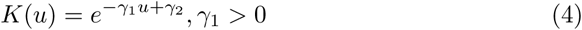

Here, *u* (0 ≤ *u* ≤ 1) represents the fraction of nucleosomes in the enhancer region that are modified with the active marks, *γ*_*i*_ are constants.

During DNA replication, markers of the two copies of H3 are randomly assigned to the two daughter cell DNA strands, each of which form a H3 dimer with a newly synthesized unmarked protein H3. Next, the nucleosome states of newly synthesized DNA are re-builded before mitosis by kinetic process of writing and erasing the marks following the enzyme-regulated biochemical reactions [28, 50, 70]. Consequently, the nucleosome states of the daughter cells depend on those of the mother cell though a random dynamics. Through a stochastic simulation based on the detail biochemical reactions of this process, we have shown that the probability of the number of marked nucleosomes of the daughter cell is approximately a conditional binormal distribution through the nucleosome state of the mother cell [28]. Thus, we made the following assumptions. Consider a DNA region enclosed in *N* nucleosomes, let *u*_*k*_ (0 ≤ *u*_*k*_≤ 1, for any *k*) the fraction of marked nucleosomes at the *k*^th^ cycle, the probability density of *u*_*k*+1_, given the value *u*_*k*_, is given by [28]

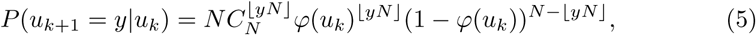

here *φ* is a predefined function so that the mean of *u*_*k*+1_, given *u*_*k*_, satisfies

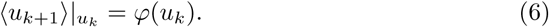

In (5), 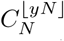 is the combination number of taken ⌊*yN*⌋ from *N*, and ⌊*yN*⌋ means the greatest integer less than or equal to *yN*. The function *φ* is crucial for defining the random inheritance of histone modification from mother cell and daughter cells. In this study, we took *φ* as linear and Hill type functions, respectively.

Finally, there are often extrinsic noise to gene expression due to environmental fluctuations. To consider the effects of extrinsic noise perturbation to the translational efficiency *λ*_*p*_, we replaced the constant *λ*_*p*_ with a log-normal distribution random process[2, 36, 56, 69].

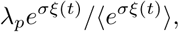

where *ξ*(*t*) is a standard white noise, *σ* > 0 is a constant to indicate the perturbation strength. Log-normal rather than normal distribution has been applied to fit the decay profile of eukaryotic mRNA [60] and many science issues [40].

In summary, we have equations for the gene expression dynamics that combine a positive feedback with histone modifications. Let *T* the duration of one cell cycle, the equation for the concentrations of mRNA *m* and protein *p* at the *k*^th^ cycle ((*k-*1)*T* < *t* < *kT*) was formulated as a random differential equation

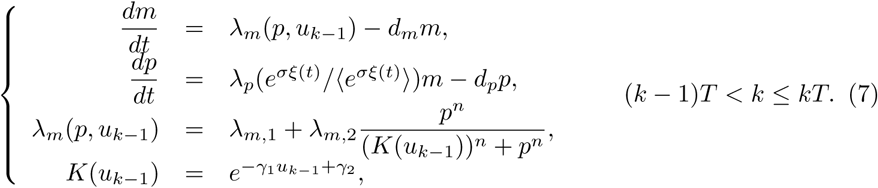

Given the initial nucleosome state *u*_0_ and the transition function *φ* in (6), the nucleosome state *u*_*k*_ at the *k*^th^ is determined iteratively by the random map in according with (5). Thus, given the initial conditions *m* and *p* at time *t* = 0 and *u*_0_ at the first cycle, and assuming that *m* and *p* are continuous cross cell cycles, the random differential equation (7) and the stochastic process (5) together define a random dynamics of the long-term gene expression cross many cell cycles.

In this study, we selected model parameter values referred to mammalian cells (human fibroblast), which are summarized in Table 1.

**Table 1.**
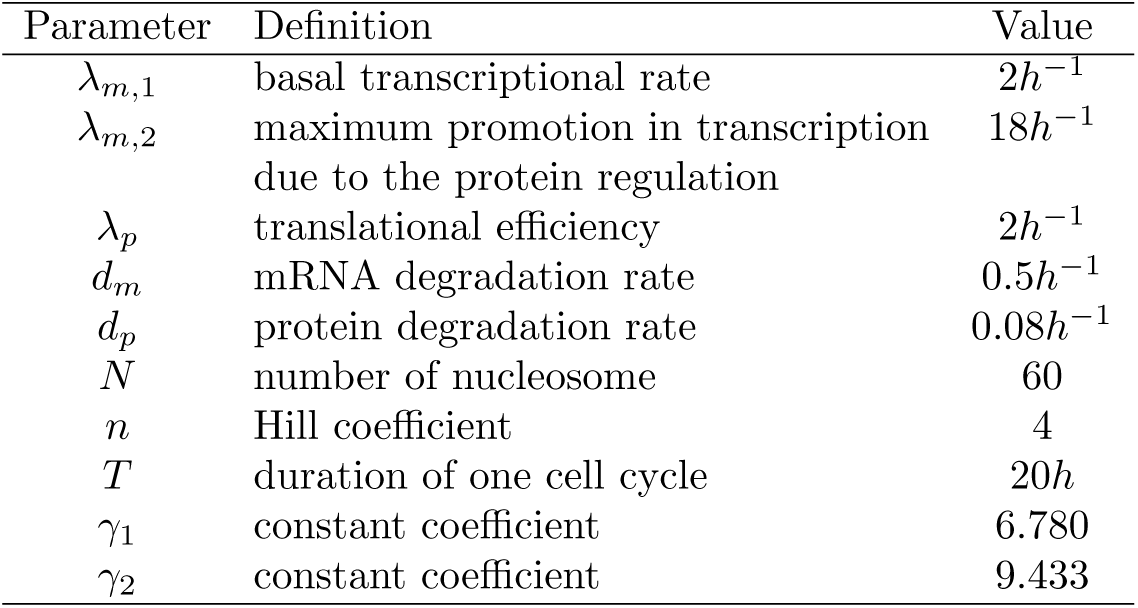
Model parameters used in simulations.

| Parameter | Definition | Value |
| --- | --- | --- |
| $\lambda_{m,1}$ | basal transcriptional rate | $2h^{-1}$ |
| $\lambda_{m,2}$ | maximum promotion in transcription due to the protein regulation | $18h^{-1}$ |
| $\lambda_p$ | translational efficiency | $2h^{-1}$ |
| $d_m$ | mRNA degradation rate | $0.5h^{-1}$ |
| $d_p$ | protein degradation rate | $0.08h^{-1}$ |
| $N$ | number of nucleosome | 60 |
| $n$ | Hill coefficient | 4 |
| $T$ | duration of one cell cycle | $20h$ |
| $\gamma_1$ | constant coefficient | 6.780 |
| $\gamma_2$ | constant coefficient | 9.433 |

## 3. Result

### 3.1. External noise-induced cell-type switch

First, we considered the system without the effect of histone modifications by assuming the nucleosome state *u* a constant over time, *i.e.*, the EC50 *K* is a constant. In this case, we have a random dynamical equation

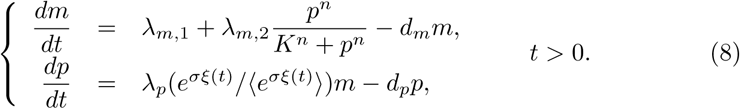

When the external noise was ignored (*σ* = 0), equation (8) describes a simple motif of self-activation, and has bistable steady states when there is cooperative binding (*n* > 1) and other parameters satisfy (Figure 2, see Appendix A for detail)

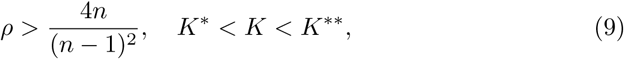

where *ρ* = *λ*_*m,*2_*/λ*_*m,*1_, and (here *α* = (*λ*_*p*_*λ*_*m,*1_)*/*(*d*_*p*_*d*_*m*_))

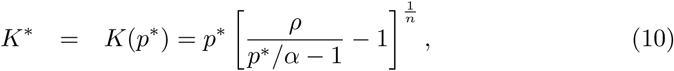

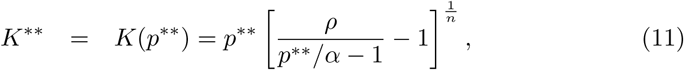

and

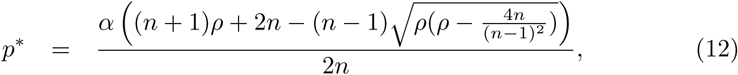

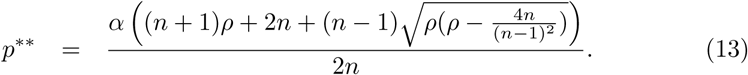

**Figure 2.**
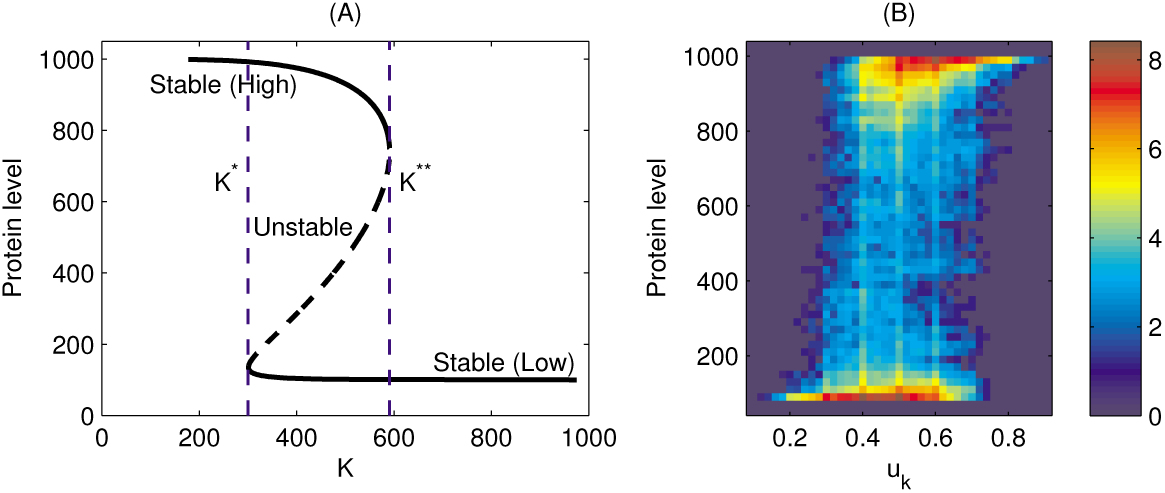
The bifurcation diagram with respect to *K* and *u*. (A) The protein level at steady state as a function of *K*. Dashed lines show the saddle-node bifurcations (*K*^*^ and *K*^**^). (B) The heat map representation of joint density distribution of the protein level and *u* for a typical trajectory without the external noise. In the simulation, *a* = 0.1, *b* = 0.8.

Here *K*^*^ and *K*^**^ are the saddle-node bifurcations, corresponding to the critical value of cell-type switches.

Since *K* depends on the nucleosome state parameter *f* through

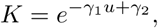

the critical values for the nucleosome state are given by

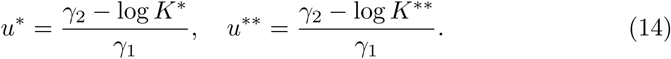

Figure 2A shows that bifurcation diagram with respect to the feedback strength *K*.

Now, we focus our discussion at the parameter region (9) so that the system is bistable; the low and high stable states mark the two distinguished cell types. While we take *K* = 450 and a suitable noise (*σ* = 1.8), the protein level switches between the two well-separated states: the low state with 0 < *p* < 500 and the high state with *p* > 500 (Fig. 3A). To further investigate how cell-type switches depend on the external noise, we fixed the parameters *λ*_*m,*1_, *λ*_*m,*2_, varied *K* over the interval *K*^*^ < *K* < *K*^**^, and increase the noise strength *σ* from 0 to 2.0. For each pair of parameters (*K, σ*), we initialized the state (both mRNA and protein concentrations) at the low level cell type, solved the equation (8) to 10^5^ cycles. We calculated the frequency (times per cycle) of cell-type switches between low and high states that is represented by dividing the number of cell-type switches by the total cell cycles. The obtained frequency is shown at Figure 3B, which indicated that noise-induced cell-type switches occur only when the noise strength *σ* is larger than 1.5. A sample dynamics of *λ*_*p*_ with *σ* = 1.5 is shown at Figure 3C, which corresponds to a large noisy effect on the parameter *λ*_*p*_. These results show that without the effects of histone modifications, cell-type switches occur only when the noise *σ* is very large.

**Figure 3.**
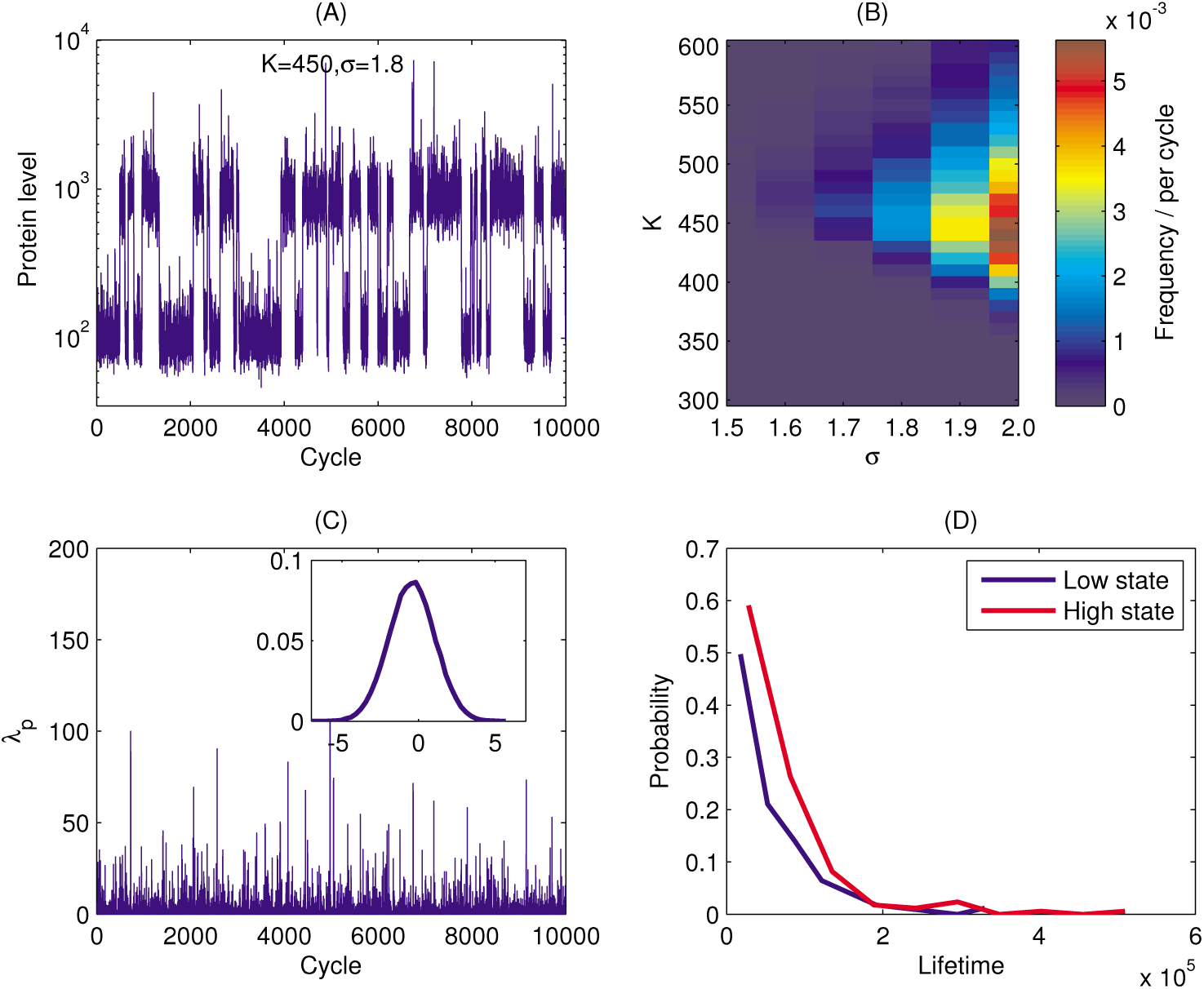
Cell-type switches induced by external noise. (A) Sample dynamics of protein level in 10^5^ cycles. In the simulation, *K* = 450, *σ* = 1.8. (B) The heat map representation of the switching frequency with respect to *K* and *σ*. (C) A sample dynamics of *λ*_*p*_ with *σ* = 1.5. Inset is the probability distribution of ln(*λ*_*p*_). (D) The probability distribution of the steady state lifetimes with correspond to the trajectory in (A).

To quantify the lifetime of the two states under noise perturbation, we defined the lifetime based on model simulation. Let the solution *p*(*t*) of a cell with initial conditions at the low state, we defined the time series {*t*_*i*_} and {*s*_*i*_} as

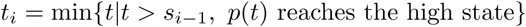

and

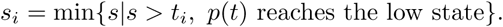

It is easy to have

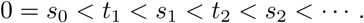

The duration time (lifetime) of the solution (cell) staying at the high state and the low state can be estimated by the two time series

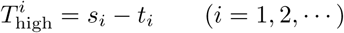

and

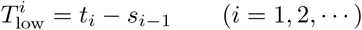

respectively. The series 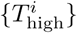 and 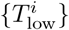 gives possible lifetimes of high and low level states, which are random numbers with exponential distribution (Fig. 3D). From Figure 3C, there is no obvious difference between the lifetime of high and low level states.

### 3.2. Stochastic histone modification-induced cell-type switch

Now, we considered the effects of random histone modification inheritance to gene expression. In our model, the state of histone modifications *u*_*k*_ stochastically change in cell cycle following the random dynamics described by (5) and (6). First, we considered a simplest case by assuming a linear function *φ*,

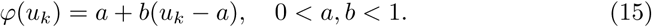

Here, the parameter *a* gives the fixed point of the iteration *u*_*k*+1_ = *φ*(*u*_*k*_). Thus, The parameters *a* and *b* are crucial for determining the random dynamics of the histone modification states {*u*_*k*_}, which is well defined by the binomial distribution (5)-(6) and (15) (Fig. 4A).

**Figure 4.**
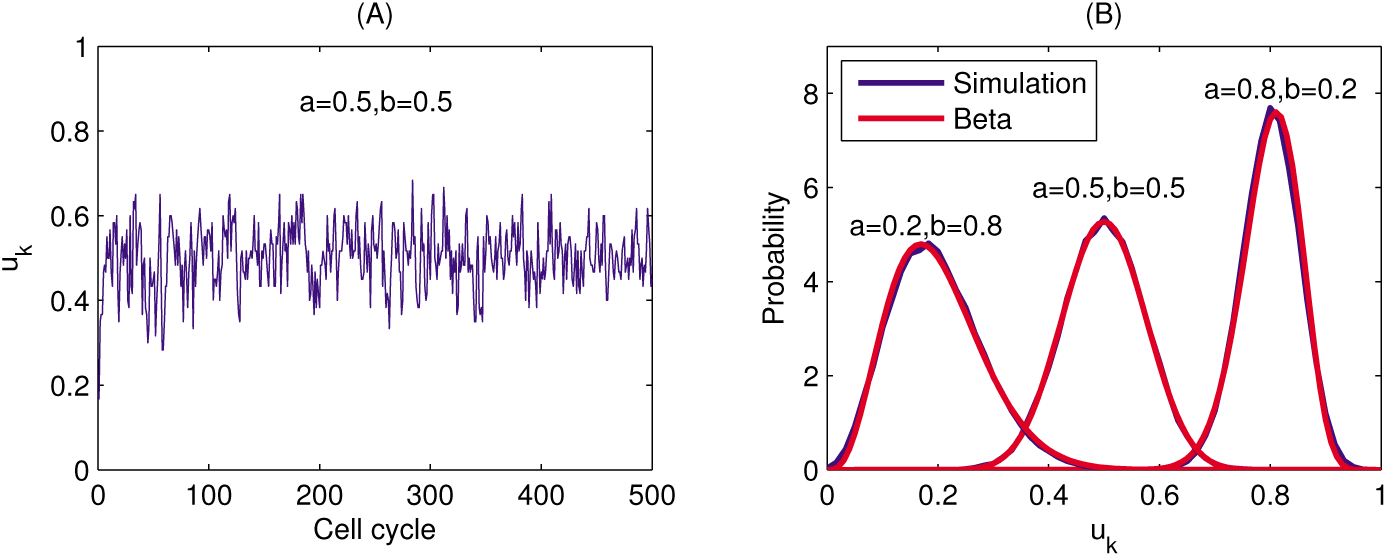
Dynamics of nucleosome states under random histone modification inheritance. (A) A sample dynamics of the nucleosome state along with cell cycles. (B) The distribution (blue) of the sequence {*u*_*k*_} with three sets of parameters of *a* and *b*, and the Beta distribution density function (red) (16) with shape parameters (*α, β*) given by (17).

The stationary distribution of *u*_*k*_ can be well described by a Beta distribution random number (Fig.4B). Specifically, a Beta distribution random number is a random number 0 ≤ *x* ≤ 1 with probability density function

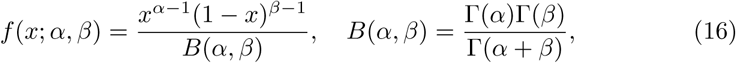

where *α* and *β* are shape parameters, and Γ(*z*) is the gamma function. The mean and variance of a Beta distribution are given by

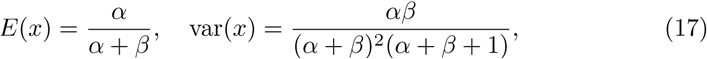

respectively. Hence, given the parameters *a* and *b* in the function *φ*, we can obtain the random dynamics *u*_*k*_ and the corresponding mean and variance, from which we can solve the Beta distribution parameters *α* and *β* from (17). Figure 4B shows the density functions of *u*_*k*_ obtained from numerical simulation and from the Beta distribution with parameters (*α, β*) from (17), which show good agreement.

In previous discussions, the EC50 parameter *K* in the transcription rate (3) is dependent on the nucleosome state *u*_*k*_ through (4). When there is no external noise (*σ* = 0), the transcription rate depends on the nucleosome state *u*_*k*_, and cell-type switch occurs when *u*_*k*_ cross the critical values *u*^*^ or *u*^**^. Figure 2B gives the distribution of protein level *p* and nucleosome state *u* along a trajectory, which show clear bistability with *u* is taking intermediate values. Based on the Beta distribution of the nucleosome state {*u*_*k*_}, the frequency of cell-type switch is dependent on the parameters *a* and *b*, which can be adjusted biologically through the regulation of the stochastic histone modification inheritance. In particularly, it is easy to induce bi-directional switch by proper selected parameters *a* and *b* in the inheritance function (15) (to be detailed below).

To examine how the frequency of cell-type switch depends on the stochastic histone modification, we fixed *σ* = 0, and varied both *a* and *b* from 0.1 to 0.9. The frequencies of cell-type switches between low and high states are shown at Figure 5A-B. The switching frequency is mainly dependent on *a*, and has particularly high frequency when *a* is closes to 0.5, regardless of the value of *b*. Hence, for a given value *b*, we can adjust the switching frequency by varying the parameter *a*, the average of nucleosome state *u*_*k*_ at the stationary state.

**Figure 5.**
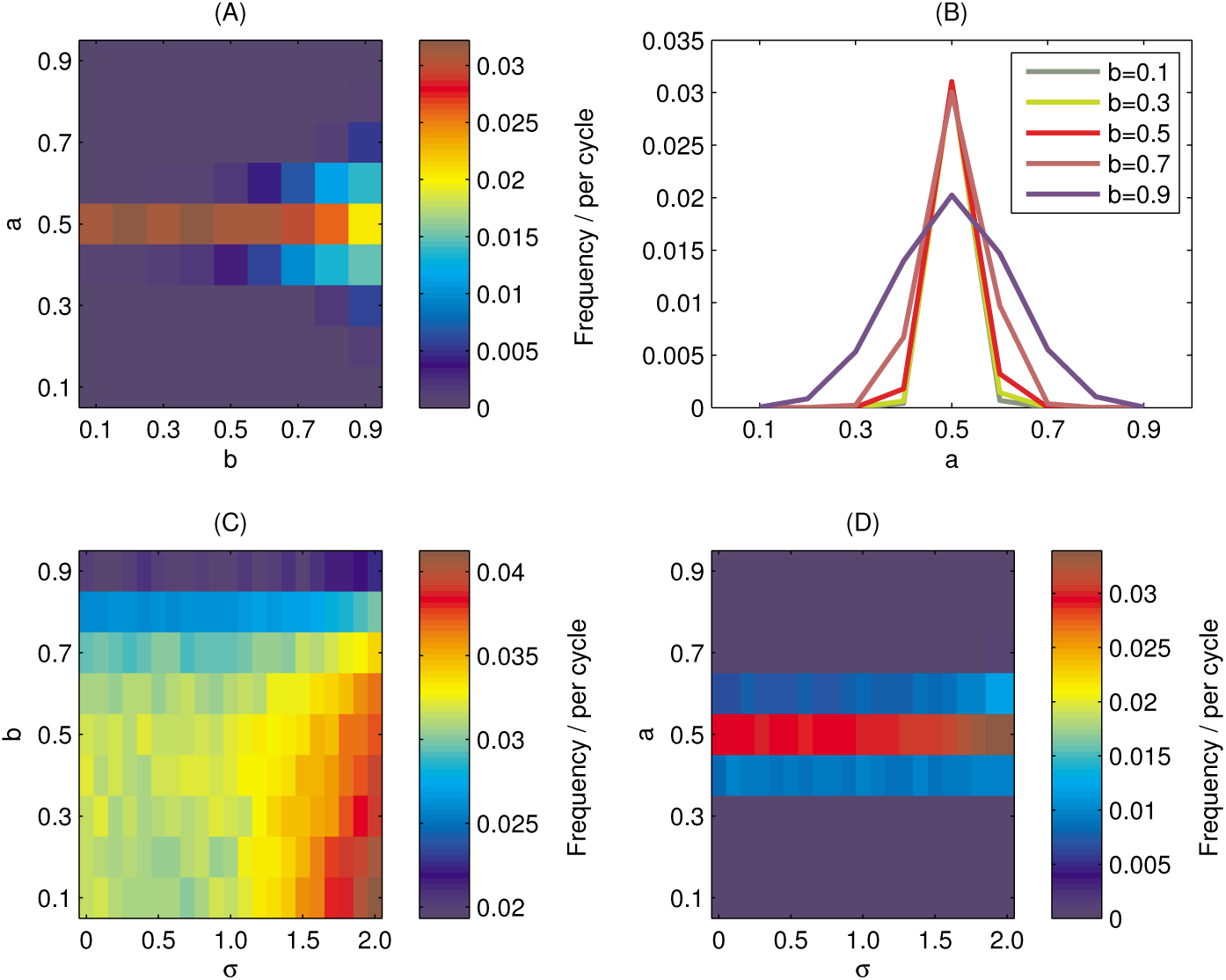
Frequencies of cell-type switches with histone modification to affect the transcription activities. (A) The heat map representation of the switching frequency with respect to *a* and *b* without the external noise. (B) The switching frequency as a function of *a* with varying parameter *b*. (C) The heat map representation of the switching frequency with respect to *b* and the noise strength *σ* (here *a* = 0.5). (D) The heat map representation of the switching frequency with respect to *a* and the noise strength *σ* (here *b* = 0.7).

Next, we study the dependence of the switching frequency with external noise *σ*. First, we fixed *a* = 0.5, varied *b* from 0.1 to 0.9, and increased *σ* from 0 to 2.0. Results show that the switching frequency is insensitive to the parameter *b* and the external noise strength *σ* unless the external noise is large (*σ* > 1.5) (Fig. 5C). Next, we fixed *b* = 0.7, varied *a* from 0.1 to 0.9, and increased *σ* from 0 to 2.0, the switching frequency is mainly determined by the parameter *a* (Fig. 5D). These results show that through the regulation of histone modifications, cell-type switches occur even when there is a weak external noise, an the cell-type switching frequencies are insensitive with the external noise strength *σ*, and are mainly determined by the random dynamics of epigenetic inheritance. We note that the parameter *a* is associated with histone modification inheritance, and hence can be regulated through related enzyme activities.

### 3.3. Cell plasticity induced by stochastic histone modification

Many systems show cell plasticity, various type cells are able to switch between different types to achieve dynamical equilibrium in response to stimulations [18, 32, 49, 54, 65, 66]. In our simple motif, cell plasticity is represented by the ability of bi-directional cell-type switches. Previous studies have shown that without the stochastic histone modification inheritance, external noise induced bi-directional cell-type switch can happen only when the noise strength is large enough (*σ* > 1.5) (also refer [37]). Here, we examined how regulations with histone modification is cable of induing the bi-directional switch and cell plasticity.

To investigate the dynamics of cell plasticity, we modified the model to include cell proliferation. During cell proliferation, each cell divides into two daughter cells, each daughter cell inherit the mRNA and protein levels from the mother cell, and the nucleosome state *u* is determined according to the distribution (5). We set the external noise strength *σ* = 0.5, and took the parameters *a* = 0.5 and *b* = 0.7, and started model simulation from a single cell with random initial condition. The distribution of protein levels in the population evolve following population doubling, and the cell population increase to 2^20^ cells in 20 cycles. Simulations show that the protein distributions approach to the same stationary distribution starting from different initial states (Fig. 6A). This result reveals cell plasticity in population dynamics based on the proposed model, quantitatively explain the phenomena of dynamic equilibrium in stem cell regenerations [5, 26, 39, 68].

**Figure 6.**
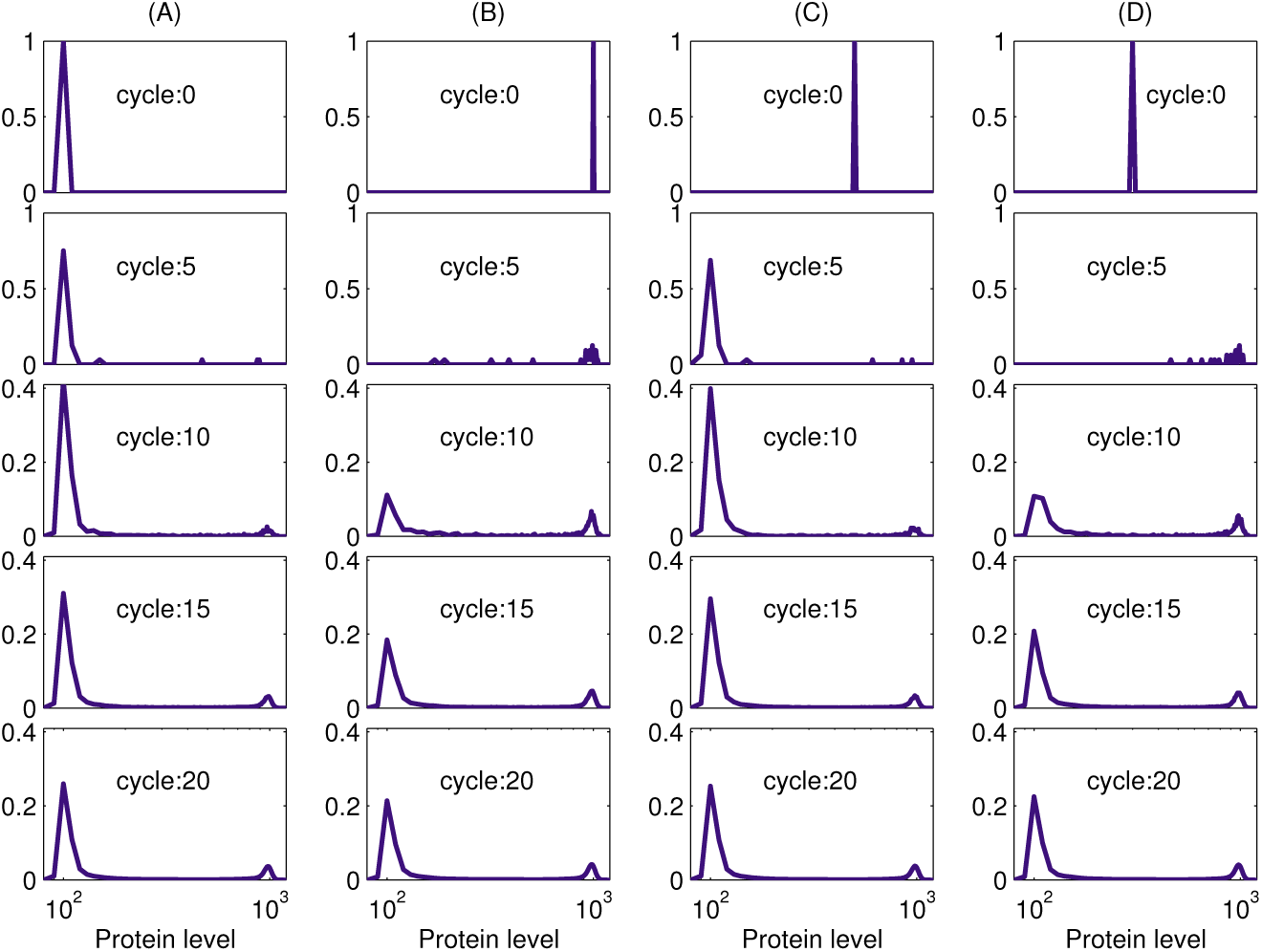
Dynamic equilibrium of cell regeneration. Figures show the probability distribution of the protein level at the end of cycle 0, 5, 10, 15, and 20. Results for 4 sample simulations ((A)-(D)) are shown, each with different initial conditions. Parameters are *a* = 0.5, *b* = 0.7.

The stationary distribution of protein levels is determined by parameters *a* and *b* for the regulations of histone modification. We fixed *b* = 0.7 and varied *a*, the protein distribution switched from the low level state to the high level state with the increasing of *a*, and display bimodal distribution when *a* is closed to 0.5 (Fig.7A). Moreover, while we fixed *a* = 0.5 and varied *b*, the stationary protein distribution is mostly unchanged with different *b* values (Fig.7B). These results show that cell plasticity can be explained by the stochastic inheritance of epigenetic states, and the epigenetic state regulation is important for determining the stationary distribution of gene expression in a colony of cell populations.

**Figure 7.**
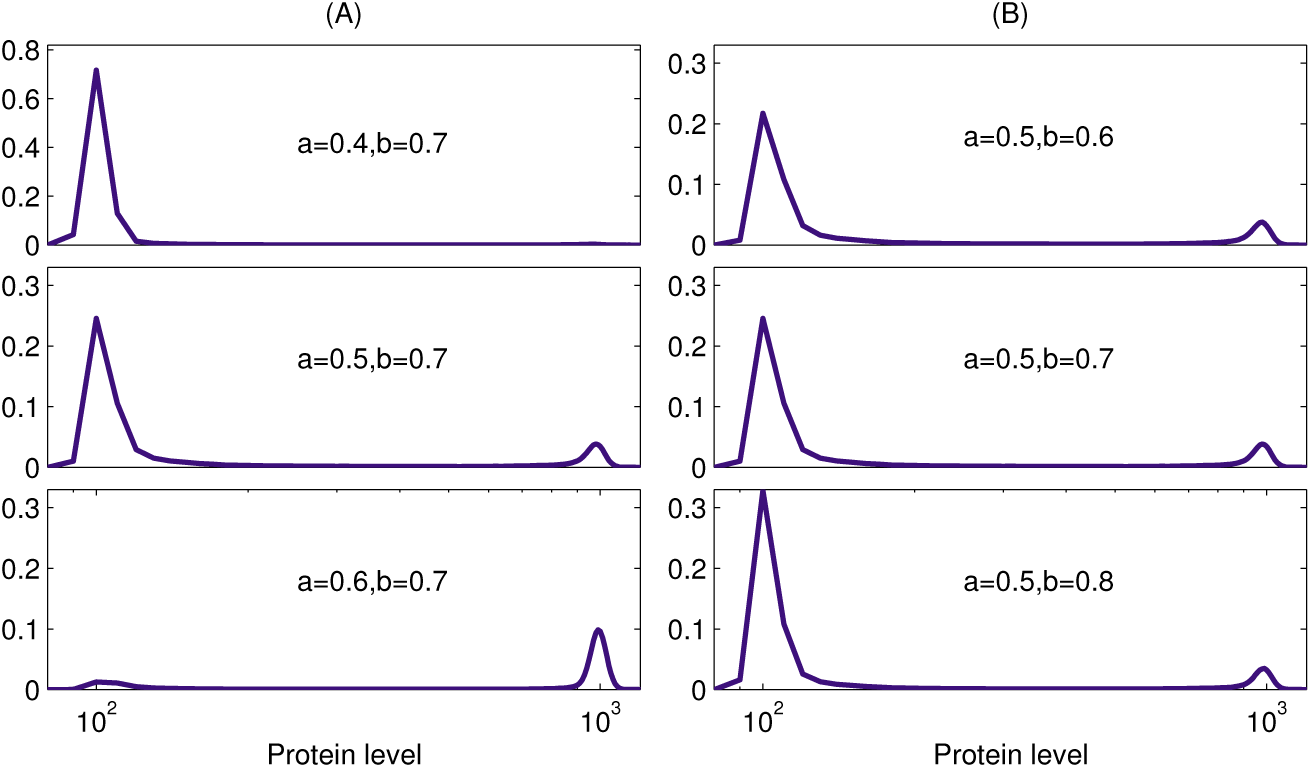
Stationary distribution of proteins in a cell colony. (A)The probability distribution of the protein level at the end of 20 cycles with fixed *b* = 0.7 and varied *a*: *a* = 0.4, *a* = 0.5, and *a* = 0.6. (B) The probability distribution of the protein level at the end of 20 cycles with fixed *a* = 0.5 and varied *b*: *b* = 0.6, *b* = 0.7, and *b* = 0.8. We take *σ* = 0.5 in simulations.

### 3.4. Waddington epigenetic landscape

To further study how stochastic histone modification affect the dynamics of cell-type switch, we compared the Waddington landscape of the phenotype development.

In the case without histone modifications, the system dynamics is described by the random dynamical equation (8). While we assumed a quasi-equilibrium to mRNA production, and the noise *σ* is small enough, we have

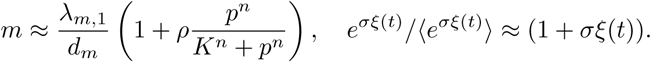

Hence, the equation (8) can be written as a stochastic differential equation

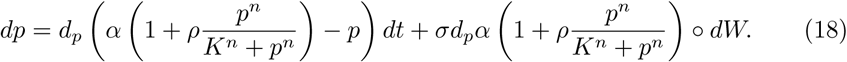

Here ∘ means the Stratonovich interpretation for the external noise[20]. Denote

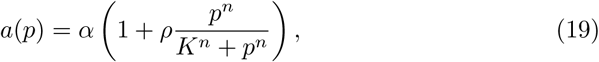

the equation (18) can be rewritten as

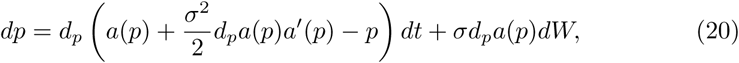

in terms of Itô interpretation [46]. Hence, the probability density of protein concentration at time *t*, *P* (*p, t*), satisfies the Fokker-Plank equation of form

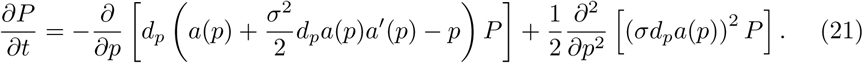

Let the stationary solution of the equation (21)

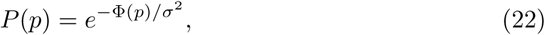

we obtain the potential landscape

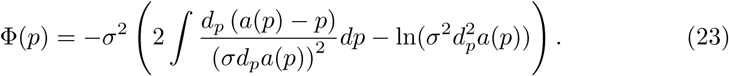

This potential landscape, which depends on model parameters, is capable of describing the stationary distribution of protein level through (22). Based on the potential Φ(*p*), varying the parameter *K* results in the switch from one local minimum to the other, which give the mechanism of cell-type switch induced by external stimulus (Fig. 8A).

**Figure 8.**
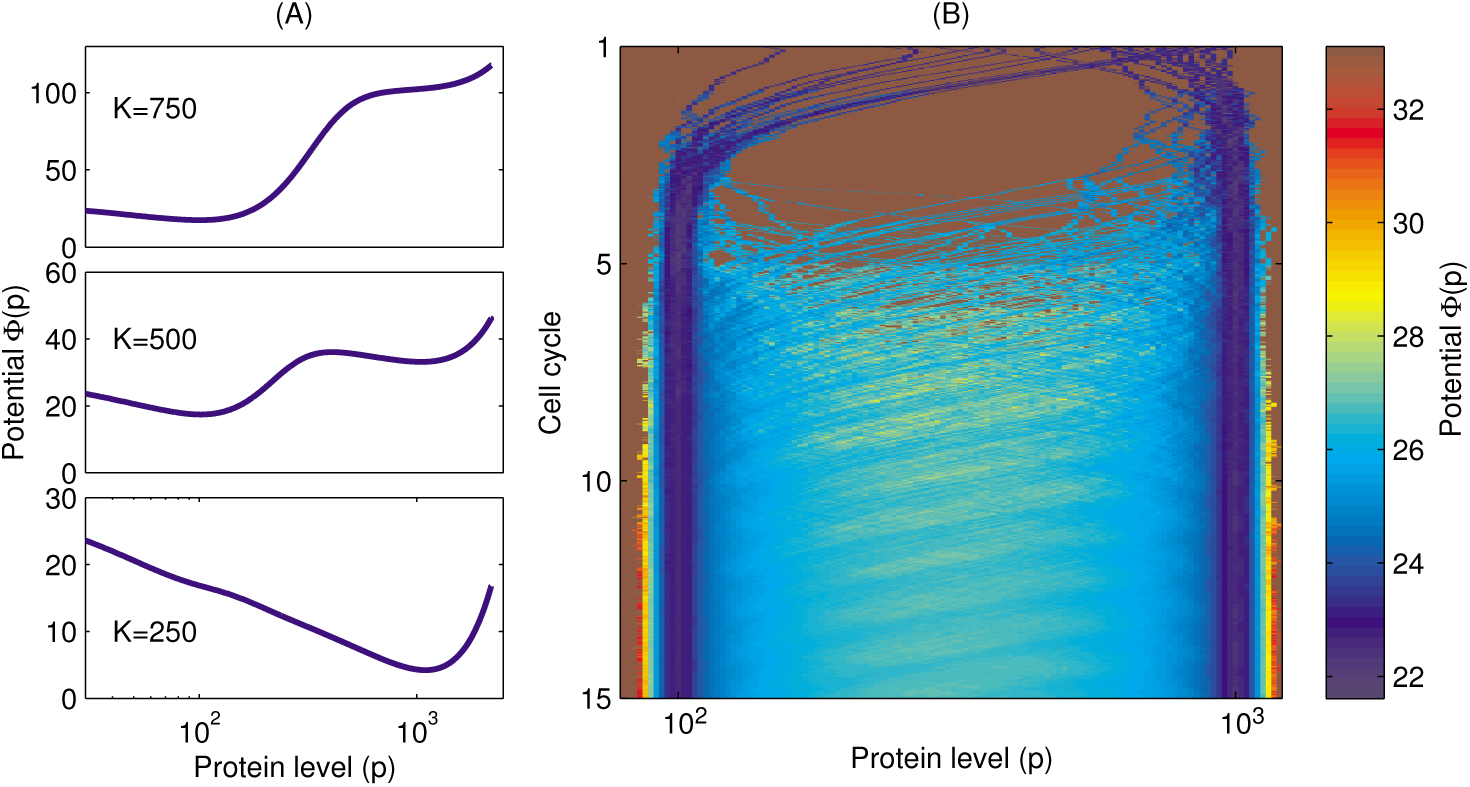
Potential landscapes. (A)The potential landscape (23) as a function of protein level with *K* = 200, *K* = 450, and *K* = 750. In the simulation, *σ* = 0.1. (B)The Waddington landscape of colony evolution from a single cells. The potential was obtained from according to simulations results of 30 independent trajectories. Parameters are *a* = 05, *b* = 0.7.

When there are stochastic histone modifications, the parameter *K* is a random dynamics during cell cycling, and hence the stationary potential landscape can not be defined in a simple way. In this case, while we consider the development of cell colony expansion, the distribution of cell state is changing over time. Hence, let *P* (*p, t*) the distribution of proteins of the cell population at time *t*, we defined the potential as Φ(*p, t*) = − ln *P* (*p, t*). Thus, the potential Φ(*p, t*) gives the evolution of epigenetic distribution, which can be considered as the Waddington epigenetic landscape of the cell population dynamics. Figure 8B shows the Waddington landscape of a colony starting from a single cell in according with the dynamics in Figure 6. From the landscape, staring from a single cell of random initial condition, the cell converges to either high or low protein level state in cycle 1; in the latter stage of colony expansion, the cells can switch between the two cell states, showing cell plasticity, and approach stable distribution of cell states.

### 3.5. Epigenetic regulation induced multistability

In many biological systems of cell-type switches, there are intermediate cell states that play important roles in mediating cell fate transitions [44]. Typical biological systems with intermediate cell states include the epithelial-to-mesenchymal transition (EMT)[63, 45], hematopoietic progenitor cell differentiation[67], and CD4^+^ T cell lineage specification. Many computational models were proposed, based on feedback regulations or noise perturbations in gene expressions, trying to understand the mechanisms and roles of intermediate cell states in cell fate decisions [71, 51].

Here, we asked whether combinations between feedback regulations and stochastic epigenetic state transition are able to induce the intermediate cell states. To this end, we assumed a cooperative effects in the histone modifications so that the transition function *φ* was taken as a nonlinear function

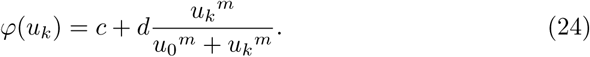

The function (24) represents a biological mechanism of positive feedback regulations in histone modifications. These type feedbacks have been considered in previous studies of nucleosome state transition [9, 59]. Under this assumption, we can adjust the parameters to obtain a dynamical process with intermediate cell states between the distinguished low and high protein level states (Fig. 9A). From Figure 9A, the cell can either switch directly between the low and high level states, or indirectly through the intermediate state, which show similar dynamics as in the intermediate state of EMT[44]. Moreover, in the simulation, the nucleosome state shows bimodal distribution with two distinguish states (Fig. 9B), however the protein level shows multi-modal distribution with three stable stale (Fig. 9C). This result suggests that the intermediate state is the consequence of the combination between regulations in the nucleosome state (*u*) and the positive feedback in gene expression. These results reveal a novel mechanism to induce the multistep transition in cell-type switches, in which the changes in epigenetic state play important roles as a buffer of phenotype changes in cell fate decision.

**Figure 9.**
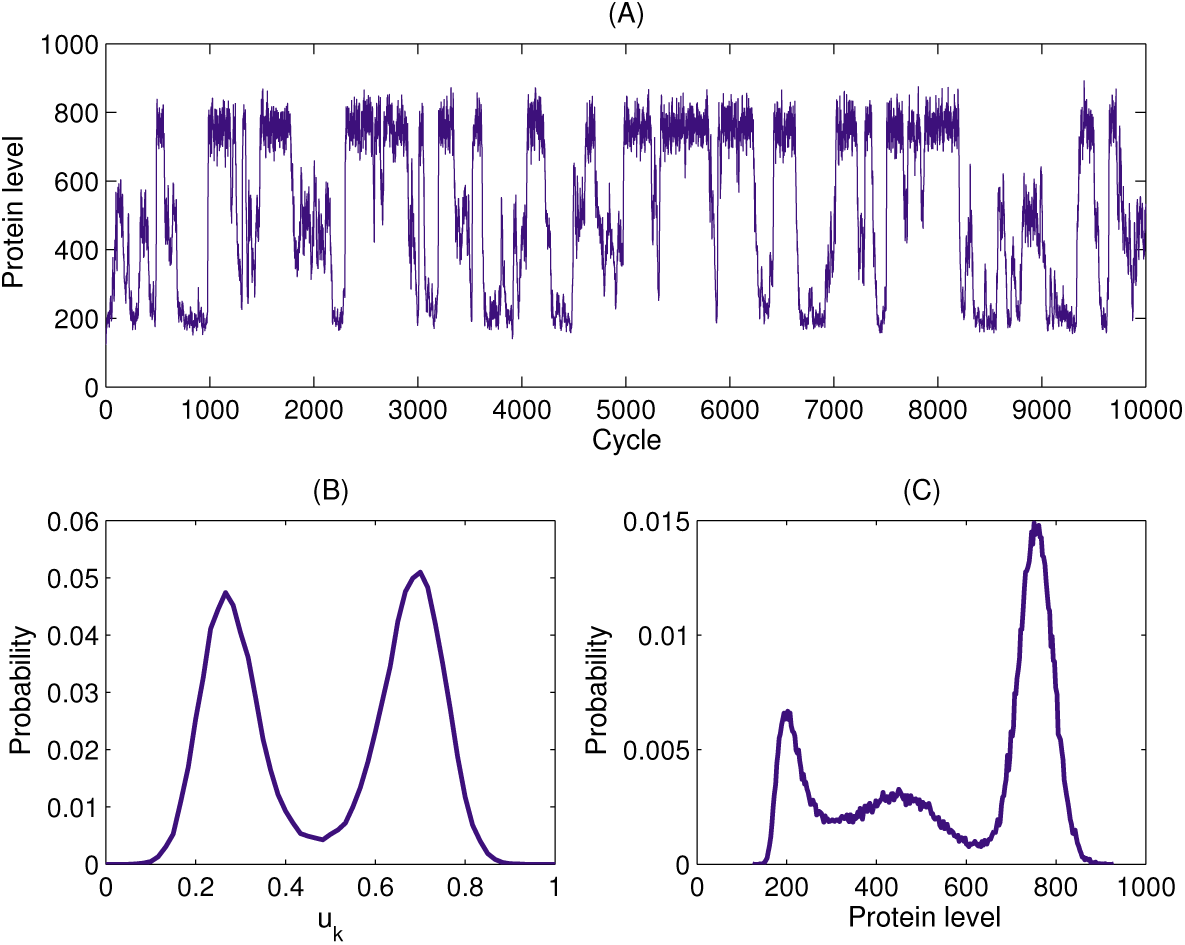
Dynamics of intermediate cell states. (A) A sample dynamics of transitions between three cell states based on the assumption (24). (B) The distribution of nucleosome state *u*_*k*_. (C) The distribution of the protein levels along the trajectory. In the simulation, we took *m* = 6, *c* = 0.25, *d* = 0.5, *u*_0_ = 0.488, *n* = 2.5, *γ*_1_ = 0.499, *γ*_2_ = 6.471, *σ* = 0.5, and the other parameters are the same as those in Table 1.

## 4. Discussion

Advances in single-cell technologies have enable us to study cell states in high resolution levels, including identification of new cell types that occupy less well-characterized roles in an atlas of cells (see the Human Cell Atlas project [53]). In particularly, the previously identified distinct cell types may connected to each other through continuous spectrum of cell-type changes [43]. These observations have led to new thoughts of stem cell differentiation and cell fate decisions from stem cell differentiation tree to complex differentiation landscapes [35]; the stem cell are flexible in space and time, and display continuum differentiation during regeneration. These new technologies have challenged the classical opinion of distinct cell states arise from dynamic attractors determined by genetic works [13, 29, 72]. In additional to the gene circuits to maintain the stable cell fates, the epigenetic states play important roles in modulating the cell plasticity along cell cycling. Alternation of histone modification or DNA methylation are often seen in stem cell differentiations and therapy-induced drug resistance [8, 11, 14].

The current study was intended to investigate how epigenetic modifications are involved in the regulation of cell-type switches. We established a simple model that couples a positive feedback with histone modification and cell cycling. Our model provides a mechanism of cell plasticity induced by the inherent random transition of histone modifications during normal cell cycle. Therewith, the flexibility of stem cells is an inherent property of cell cycling, which can be important in maintaining the heterogeneity and robust dynamics of stem cell regeneration [38]. Moreover, we found that coupling the regulations between histone modification and positive feedback gene circuit can result in the intermediate cell states that play essential role in EMT. Extension of the proposed simple model may provide insights to the future study of how inherent epigenetic regulations work together with the complex gene circuits to modulate the process of cell-type switches, as well as the understanding of drug resistance in cancer therapy.

### Appendix A. Bistability analysis

Consider the deterministic differential equation

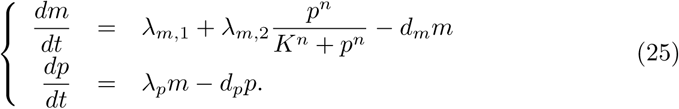

The steady state is given by the equation

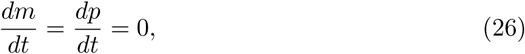

which yields

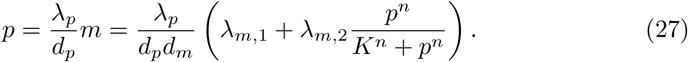

The equation (27) has multiple roots when *n* > 1 and suitable parameter values *K*. To obtain the conditions for the multiple roots, we solved *K* from (27), which gives

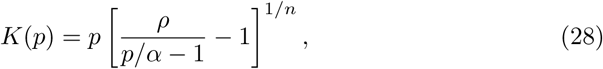

where

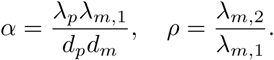

It is easy to verify that any root *p* of (27) satisfies

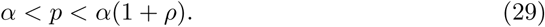

Hence, we limited our discussion to the condition (29). The critical point *K*^*^ for bistability corresponds to the value *K* = *K*^*^ so that

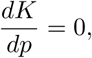

which gives

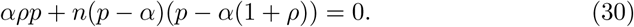

Thus, it is easy to have the following: If

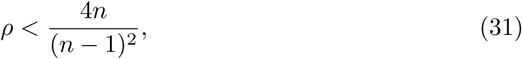

*K*(*p*) is monotonously decrease with *p*, and (27) has a unique solution; if

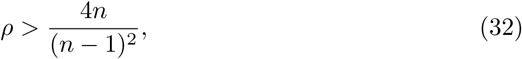

the derivative 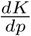 has two roots at

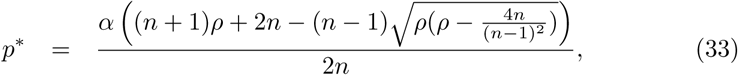

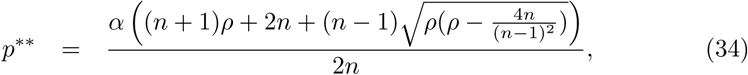

and *α < p*^*^*< p*^**^*< α*(1 + *ρ*). Thus, the function *K*(*p*) decreases when *α < p < p*^*^, increases when *p*^*^*< p < p*^**^, and decreases when *p*^**^*< p < α*(1 + *ρ*). Therewith, the equation (25) has two saddle-nodes

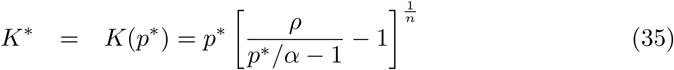

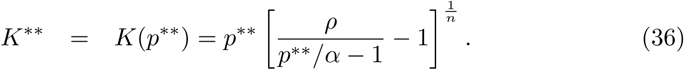

The above equations give the explicit formulation of the saddle-nodes, and (25) is bistable when *n* > 1 and

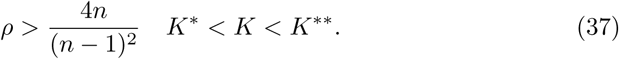

Received xxxx 20xx; revised xxxx 20xx.

*E-mail address*

*E-mail address*

